# How transposons drive evolution of virulence in a fungal pathogen

**DOI:** 10.1101/038315

**Authors:** Luigi Faino, Michael F Seidl, Xiaoqian Shi-Kunne, Marc Pauper, Grardy CM van den Berg, Alexander HJ Wittenberg, Bart PHJ Thomma

## Abstract

Genomic plasticity enables adaptation to changing environments, which is especially relevant for pathogens that engage in arms races with their hosts. In many pathogens, genes mediating aggressiveness cluster in highly variable, transposon-rich, physically distinct genomic compartments. However, understanding of the evolution of these compartments, and the role of transposons therein, remains limited. We now show that transposons are the major driving force for adaptive genome evolution in the fungal plant pathogen Verticillium dahliae. Highly variable genomic regions evolved by frequent segmental duplications mediated by erroneous homologous recombination, often utilizing transposons, leading to genetic material that is free to diverge. Intriguingly, the duplicated regions are enriched in active transposons that further contribute to local genome plasticity. Thus, we provide evidence for genome shaping by transposons, both in an active and passive manner, which impacts the evolution of pathogen aggressiveness.

## Introduction

Genomic plasticity enables organisms to adapt to environmental changes and occupy novel niches. While such adaptation occurs in any organism, it is particularly relevant for pathogens that engage in co-evolutionary arms races with their hosts (Raffaele and Kamoun 2012; Seidl and Thomma 2014; Dong et al. 2015). In these interactions, hosts utilize their surveillance system to detect invaders and mount appropriate defenses, which involves detection of invasion patterns by immune receptors, while pathogens secrete so-called effector molecules that support host colonization to counteract these immune responses (Rovenich et al. 2014; Cook et al. 2015). The tight interaction, in which the microbe tries to establish a symbiosis that the host tries to prevent, exerts strong selection pressure on both partners and incites rapid genomic diversification (McDonald and Linde 2002).

Sexual reproduction is an important mechanism to establish genotypic diversity by combining genetic information of two parental lineages, followed by meiotic recombination leading to novel combinations of existing alleles (McDonald and Linde 2002; de Visser and Elena 2007; Heitman 2010; Speijer et al. 2015). However, not all eukaryotes regularly reproduce sexually, including many fungal phyla that are thought to reproduce mainly asexually (Heitman et al. 2007; Flot et al. 2013). However, also in such asexual organisms adaptive genome evolution occurs, which is mediated by various mechanisms ranging from single-nucleotide polymorphisms to large-scale structural variations that can affect chromosomal shape, organization and gene content (Seidl and Thomma 2014). *Verticillium dahliae* is a soil-borne fungal pathogen that infects susceptible hosts through their roots and colonizes the water-conducting xylem vessels, leading to vascular wilt disease (Fradin and Thomma 2006). Despite its presumed asexual nature, *V. dahliae* is a highly successful pathogen that causes disease on hundreds of plant hosts (Fradin and Thomma 2006; Klosterman et al. 2009; Inderbitzin and Subbarao 2014). Verticillium wilt diseases are difficult to control due to the long viability of fungal resting structures in the soil, the broad host range of the pathogen, and inability of fungicides to eliminate the pathogen from infected xylem tissues. Moreover, little genetic resistance is available (Fradin and Thomma 2006). Thus, novel methods for disease control are urgently required, which requires increased understanding of the biology of the fungus in its interaction with host plants (Fradin and Thomma 2006).

Using comparative genomics, we recently identified genomic rearrangements in *V. dahliae* that lead to extensive chromosomal length polymorphisms between strains (de Jonge et al. 2013). Moreover, the genomes of *V. dahliae* strains were found to contain highly dynamic, repeat-rich regions that are only present in subsets of the *V. dahliae* population, and thus have been referred to as lineage-specific (LS) (Klosterman et al. 2011; de Jonge et al. 2013). Intriguingly, LS regions are enriched for *in* planta-induced effector genes that contribute to fungal virulence (de Jonge et al. 2013). Therefore, it was hypothesized that elevated genomic plasticity in LS regions contributes to diversification of effector repertoires of *V. dahliae* lineages and thus mediates their aggressiveness (de Jonge et al. 2013; Seidl and Thomma 2014). In a similar fashion, many filamentous pathogens evolved so-called ‘two-speed’ genomes with gene-rich, repeat-poor genomic compartments that contain core genes that mediate general physiology and evolve slowly, whereas plastic, gene-poor and repeat-rich compartments are enriched in effector genes that mediate aggressiveness in interactions with host plants and evolve relatively quickly (Raffaele and Kamoun 2012).

Plastic, fast-evolving genomic compartments in plant and animal pathogen genomes concern particular regions that are either embedded within the core chromosomes or reside on conditionally dispensable chromosomes (Thon et al. 2006; Fedorova et al. 2008; Haas et al. 2009; Ma et al. 2010; Goodwin et al. 2011; Klosterman et al. 2011; Rouxel et al. 2011; de Jonge et al. 2013). Irrespective of the type of organization of the ‘two-speed’ genome, the fast-evolving compartment is generally enriched for transposable elements (TEs). These elements have been hypothesized to actively promote genomic changes by causing DNA breaks during excision, or by acting as a substrate for rearrangement (Seidl and Thomma 2014). Nevertheless, the exact role of TEs in the evolution of effector genes currently remains unknown, as well as the molecular mechanisms and the evolutionary trajectories that govern genome plasticity (Dong et al. 2015). Thus far, in-depth studies of the precise roles of TEs were hampered by fragmented pathogen genome assemblies caused by limitations in sequencing technologies, leading to poor assemblies of repeat-rich areas particularly (Faino et al. 2015; Thomma et al. 2015). We recently obtained gapless *V. dahliae* whole-genome assemblies using a combination of long-read sequencing and optical mapping (Faino et al. 2015). Here, we exploit these novel assemblies for in-depth investigation into the molecular mechanisms that are responsible for genomic variability, and are instrumental for adaptive genome evolution in the plant pathogen *V. dahliae.*

## Results

### *Genomic rearrangements in* Verticillium dahliae *are associated with extensive sequence similarity*

To investigate the molecular mechanisms that underlie genomic rearrangements in *V. dahliae,* we performed whole-genome alignments between strains JR2 and VdLs17 (Faino et al. 2015), revealing 24 synteny disruptions (Fig.1A; Supplemental Fig. 1). To further increase the resolution of these synteny breakpoints, we subsequently aligned long (average ∽9 kb) sequencing reads derived from strain VdLs17 (Faino et al. 2015) to the completed, gapless assembly of *V. dahliae* stain JR2, and reads of which the two sides aligned to two distinct genomic locations were used to determine breakpoints at high-resolution (Supplemental Fig. 2). After manual refinement, this procedure yielded 19 large-scale chromosomal rearrangements, 13 of which concern inter-chromosomal translocations and 6 concern intra-chromosomal inversions (Fig. 1; Table 1). Of these 19 rearrangements, seven were confined to regions smaller than 100 bp (Table 1). The remaining 12 genomic rearrangements, in particular intra-chromosomal inversions, concern to larger genomic regions that could not be further refined due to the presence of strain-specific and/or repeat-rich regions (Fig. 1B).

**Table 1.**
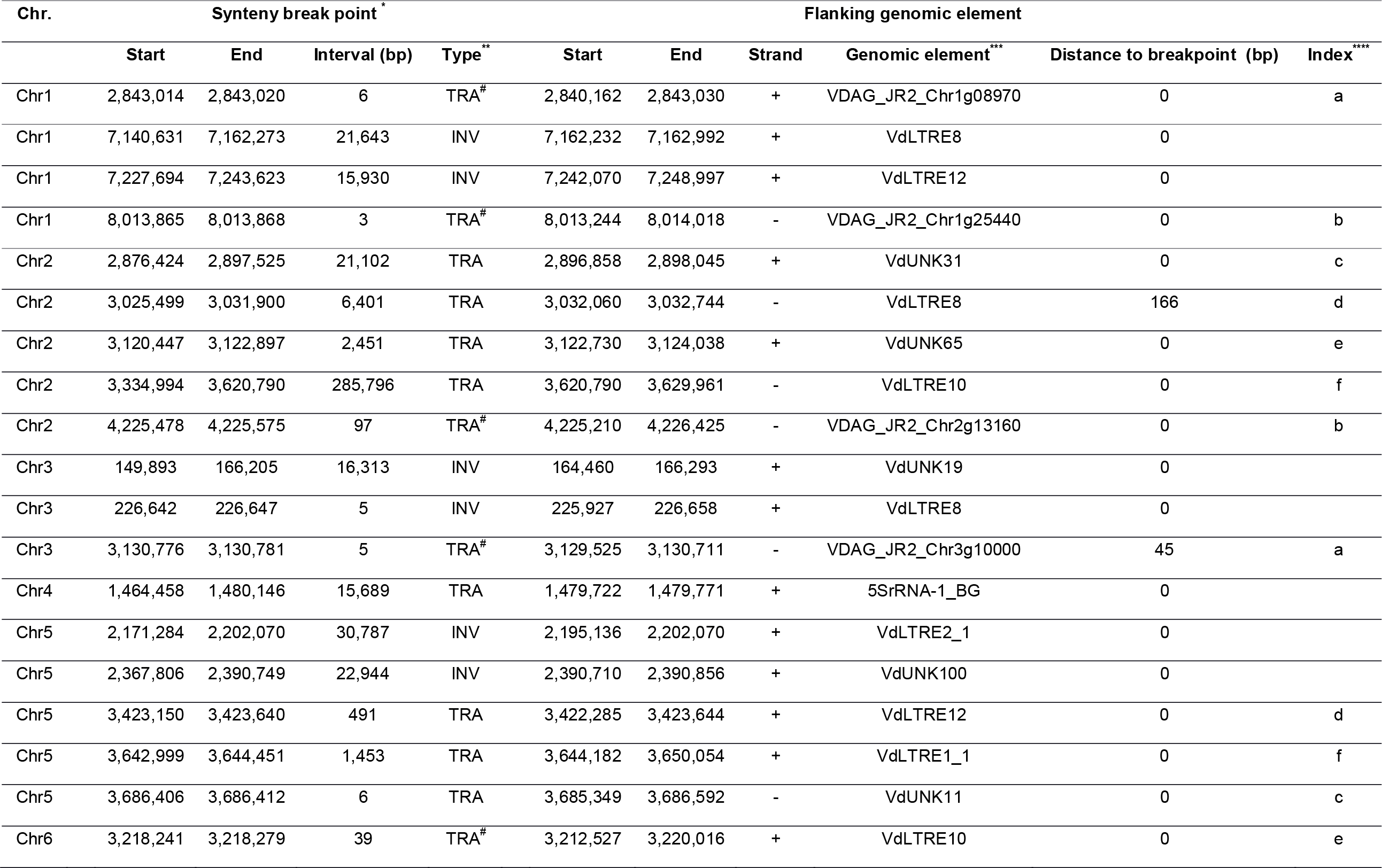
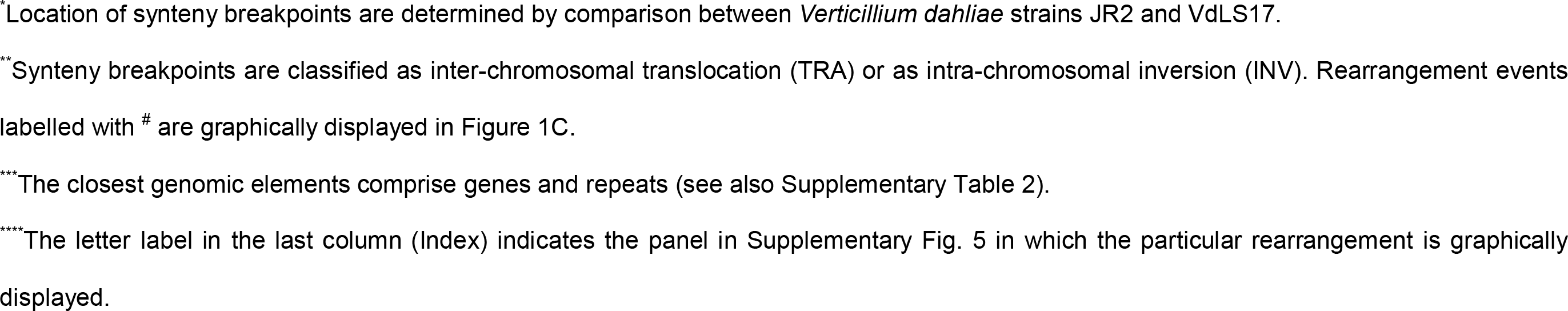
Genomic location of synteny breakpoints identified in *Verticillium dahliae* strain JR2 and the closest flanking genomic element.

**Figure 1.**
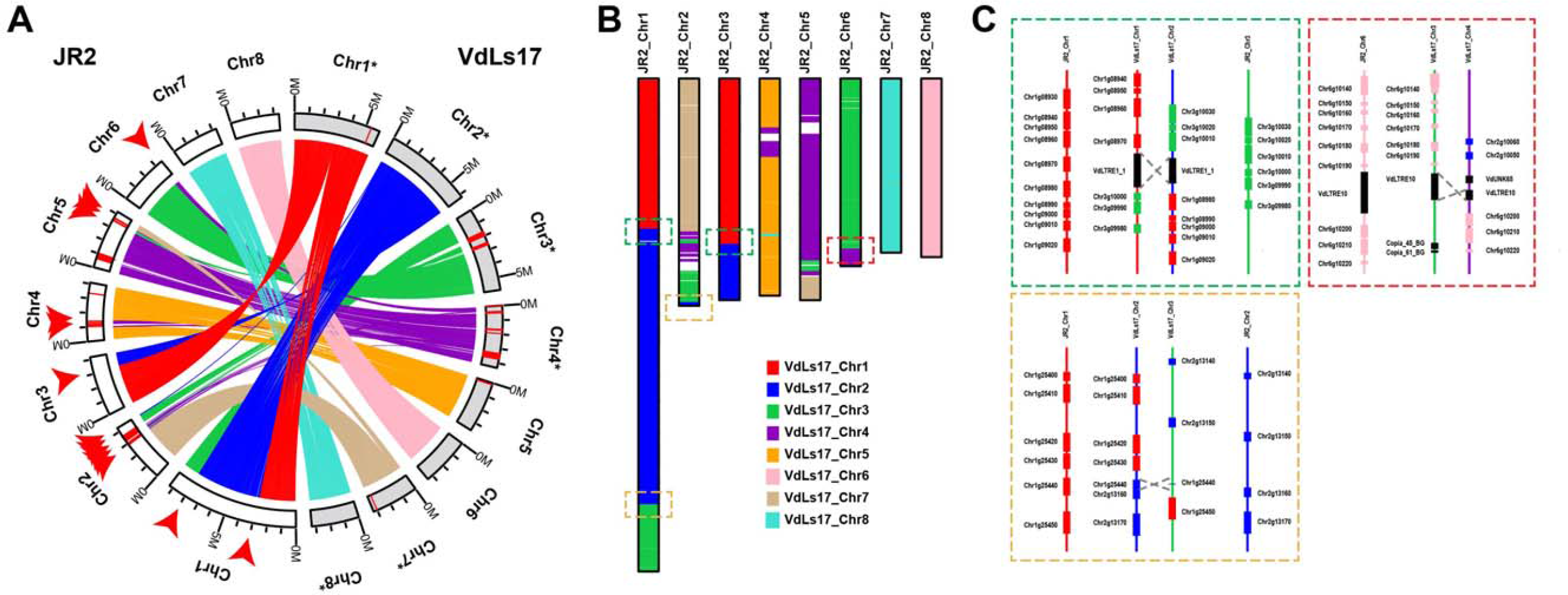
Extensive rearrangements in *Verticillium dahliae* genomes are mediated by repetitive elements. (A) Syntenic regions, indicated by ribbons, between chromosomes of the two highly similar *V. dahliae* strains JR2 (chromosomes displayed in white) and VdLs17 (chromosomes displayed in grey) reveal multiple synteny breakpoints caused by inter-chromosomal rearrangements, highlighted by red arrows for the JR2 genome. Red bars on the chromosomes indicate lineage-specific genomic regions (LS) that lack synteny in the other strain. To facilitate visibility, some chromosomes of *V. dahliae* strain VdLs17 have been reversed and complemented (indicated by asterisks). (B) Synteny blocks displayed on the chromosomes of *V. dahliae* strain JR2 colored based on synteny with VdLs17 chromosomes. LS regions are shown in white and indicate loss of synteny. (C) Detailed view of the genomic regions surrounding five synteny breakpoints that are highlighted by boxes in (b). Rearrangements over short homologous regions such as repetitive elements (black boxes) or genes (colored boxes) resulted in inter-chromosomal rearrangements (translocations). *V. dahliae* strain VdLs17 genes were inferred by mapping of the *V. dahliae* strain JR2 genes to the genome assembly of *V. dahliae* strain VdLs17. Dashed grey lines indicate rearrangement sites.

We subsequently assessed the occurrence of the 13 inter-chromosomal rearrangements in nine other *V. dahliae* strains by querying paired-end reads derived from these genomes for pairs of discordantly mapped reads (i.e. both reads fail to map at the expected distance and/or location) when mapped onto the assembly of *V. dahliae* strain JR2 (Supplemental Fig. 3). This analysis revealed distinct rearrangement patterns among various strains (Supplemental Fig. 3B). For example, while some synteny breakpoints identified in *V. dahliae* strain JR2 are either specific to *V. dahliae* strain VdLs7 (Chr1: 2,843,014-2,843,020; Supplemental Fig. 4A) or common to all other *V. dahliae* strains (Chr1: 8,013,865-8,013,868; Supplemental Fig. 4B), some are observed in only a subset of *V. dahliae* strains (Chr3: 3,130,776-3,130,781; Supplemental Fig. 4C). Thus, genomic rearrangements commonly occur in different *V. dahliae* strains, suggesting a common mechanism driving their formation.

Genomic rearrangements are generally caused by double-strand DNA breakages followed by unfaithful repair by DNA repair mechanisms that utilize homologous sequences to repair such breaks (Hedges and Deininger 2007; Krejci et al. 2012; Seidl and Thomma 2014). Repetitive genomic elements, such as transposable elements (TEs), may give rise to genomic rearrangements by providing an ectopic substrate that interferes with faithful repair of the original break (Hedges and Deininger 2007; Seidl and Thomma 2014). Notably, out of 19 chromosomal rearrangements identified in *V. dahliae* strain JR2, 15 co-localize with a repetitive element (Table 1; Supplemental Table 1). However, even though genomic rearrangements in *V. dahliae* were previously associated with LTR-type retrotransposons (de Jonge et al. 2013), we were neither able to confirm this association nor systematically associate these structural variations to any other class of repetitive elements when using the completely assembled genomes of *V. dahliae* strains JR2 and VdLs17 (Faino et al. 2015) (Table 1). To understand the mechanisms that contributed to their formation, we studied the 13 inter-chromosomal rearrangements between *V. dahliae* strain JR2 and VdLs17 in more detail. Of these 13 rearrangements, 12 could be reconstructed in detail (Fig. 1B; Table 1; Supplemental Fig. 5). Eight of these 12 syntenic breakpoints occur over highly similar TEs in both strains, albeit that they belong to different TE families (Fig. 1C; Table 1; Supplemental Fig. 5). Closer inspection of the four remaining breakpoints revealed a TE at two of the breakpoints in *V. dahliae* strain VdLs17, while this TE was absent at those breakpoints in the JR2 strain. For the final two breakpoints no association to a TE was found in either strain, but extended sequence similarity surrounding the rearrangement site was identified (Fig. 1C). Therefore, we conclude that not necessarily TEs or their activity, but rather stretches of sequence similarity are associated with ectopic chromosomal rearrangements in *V. dahliae,* likely mediating unfaithful homology-based DNA repair. Since TEs are more abundant compared to other (types of) sequences, these are more likely to become substrate for double-strand repair pathways.

### *Lineage-specific genomic regions in* Verticillium dahliae *evolved by segmental genomic duplications*

Chromosomal rearrangements can lead to ectopic insertions, duplications or deletions of genomic material, which play relevant roles in pathogen evolution during the arms race with host plants (Seidl and Thomma 2014). Whole-genome alignments between *V. dahliae* strains JR2 and VdLs17 revealed four large (>10 kb) repeat-rich genomic regions that do not display synteny between the strains (Supplemental Fig. 6; Supplemental Table 2), suggesting that at least part of the genomic material is absent in the respective other *V. dahliae* strain and indicating extensive differential gene loss at these specific genomic locations. Moreover, mapping of reads from nine additional strains (de Jonge et al. 2012) onto the JR2 and VdLs17 assemblies revealed absence of read coverage mainly at their respective LS regions (Supplemental Fig. 7). Notably, several inter-chromosomal rearrangements co-localize with these LS regions (Fig. 1A), suggesting that genomic rearrangements contribute to the formation of LS regions.

Chromosomal rearrangements not only lead to deletion of genetic material, e.g. in LS regions (Fig. 1A), but also foster genomic duplications (Seidl and Thomma 2014). Even though duplicated genes have previously been observed in LS regions of *V. dahliae* (Klosterman et al. 2011), the extent of such duplications and their role in the evolution of LS regions remains unknown. To determine the extent of segmental duplications in LS regions, we employed two approaches. First, we performed whole-genome nucleotide alignments *of V. dahliae* strain JR2 to itself to identify highly similar (>80% identity), large-scale duplication events (Fig. 2A). Unanticipated, the vast majority of highly similar large-scale duplications occurs within LS regions, while only very few of such duplications occur outside these regions (Fig. 2A). Second, we performed homology detection between protein-coding genes in *V. dahliae* strain JR2, establishing a set of ∽1,000 paralogous sequences (Fig. 2B). Notably, 40% of the 418 genes located in LS regions have a paralog (Fig. 2B), which is a 4.5x enrichment when compared to the core genome where only 7% of the ∽11,000 genes has a paralog (hypergeometic test; p = 1.31*10^-69^). Therefore, duplications of genomic material are instrumental for the constitution of LS regions in *V. dahliae* (Fig. 2B).

The high level of similarity between sequences located at LS regions (Fig. 2A) suggests that the duplications occurred rather recently. To firmly establish when these duplications occurred during the evolution of *V. dahliae,* we used the rate of synonymous substitutions per synonymous site (Ks) between paralogous gene pairs as a proxy for time since these sequences diverged (Fig. 2C). While the Ks distribution of paralogous pairs located in the core genome displays a single peak, indicating a single and distinct period in which the majority of these duplications occurred, the distribution of paralogous pairs where at least one gene is located in the LS regions displays two distinct peaks (Fig. 2C). Notably, the older of the two peaks coincides with the peak observed for the core paralogs (Fig. 2C), indicating that the expansion of core genes and a subset of genes in LS regions occurred in the same period. The additional peak points towards duplications that occurred more recently (Fig. 2C). To place these periods in relation to speciation events, we estimated Ks distributions for orthologous gene pairs between *V. dahliae* strain JR2 and a number of closely related fungi from the taxonomic class of Hypocreomycetidae (Fig. 2D). Within this group of close relatives, the tomato wilt pathogen *Fusarium oxysporum* f.sp. *lycopersici* was the first one to diverge from the last common ancestor, while the most recent split was the divergence of *V. dahliae* strains JR2 and VdLs17 (Fig. 2D). The first duplication period affected both the core genome and LS compartments, and occurred after the divergence of *F. oxysporum* f.sp. *lycopersici,* but before the divergence of *Colletotrichum higginsianum,* the causal agent of anthracnose leaf-spot disease of crucifers (Fig. 2D). Notably, all fungi that diverged after speciation of *F. oxysporum* f.sp. *lycopersici* display a peak in the Ks distribution for paralogous gene pairs at approximately the same period, although *A. alcalophilium* shows a less pronounced peak (Fig. 2D). The second duplication event in *V. dahliae* strain JR2 that specifically concerned genes located at LS regions occurred much more recently, after the speciation of *Verticillium alfalfae* (Fig. 2D). As it was previously shown that genes in the LS regions are particularly relevant for pathogen aggressiveness in *V. dahliae* (de Jonge et al. 2013), this suggests that recent gene duplications are contributing to the evolution of aggressiveness.

The recent duplications that affected LS regions have generated raw genetic material that can be subject to subsequent rapid evolutionary diversification, leading to novel or altered gene functionality, but can also be subject to differential loss of one of the duplicated gene copies (Fig. 3A; Supplemental Fig. 8). In general, LS regions in *V. dahliae* strain JR2 display considerable gene loss, since for ∽100 of the ∽400 genes located in LS regions no ortholog could be detected in *V. dahliae* strain VdLs17. For example, a repeat-rich, ∽100 kb LS region that contains ∽20 genes that duplicated on chromosome 2 displays loss of multiple genes (Fig. 3A; Supplemental Fig. 8), a process that is similarly observed in VdLs17. Thus, differential gene loss significantly contributed to the diversification of LS regions. To determine if LS regions display signs of increased gene diversification and selection pressure acting on protein-coding genes, we used the rate of nonsynonymous substitutions per nonsynonymous site (Ka) as well as the ratio of Ka to Ks values calculated between orthologous gene pairs of *V. dahliae* strain JR2 and the closest related *Verticillium* species, *V. alfalfae,* as a proxy. Genes located at LS regions display moderately increased Ka values, as well as Ka/Ks values, when compared to genes residing in the core genome (median of 0.05 compared to 0.02, and 0.37 compared to 0.17, respectively). While these indicate accelerated sequence divergence of genes located within LS regions, the moderate differences also corroborate previous results (de Jonge et al. 2013; Seidl et al. 2015), suggesting that sequence divergence only plays a minor role in *V. dahliae* genome evolution.

### *The* Ave1 *effector gene is located in a highly dynamic genomic region*

As shown above, highly dynamic LS regions are characterized by frequent gene duplications (Fig. 2), and by differential gene loss (Fig. 3; Supplemental Fig. 8). Moreover, effector genes located in LS regions play decisive roles in pathogen-host interactions (de Jonge et al. 2013). For example, Ave1 is an important LS effector that determines *V. dahliae* aggressiveness on various host plants (de Jonge et al. 2012). As expected in a co-evolutionary arms race (Thomma et al. 2011; Cook et al. 2015), host recognition of this effector evolved as tomato plants that carry the immune receptor Ve1 recognize this effector to establish immunity (Fradin et al. 2009; de Jonge et al. 2012). Consequently, race 2 strains of *V. dahliae* lost *Ave1,* thus evading recognition by Ve1, leading to the capacity to infect *Ve1* tomato plants.

In *V. dahliae* strain JR2, *Ave1* is embedded in a gene-sparse and repeat-rich LS region on chromosome 5 (Fig. 3B-E; ∽550,000-1,050,000). Notably, the average number of single nucleotide polymorphisms (SNPs) inferred from three race 1 and eight race 2 strains of *V. dahliae* is significantly reduced in the area surrounding the *Ave1* locus (between 680,000 and 720,000) when compared with the surrounding genomic regions (Fig. 3B), but also compared to genome-wide SNP levels in LS and core regions (Supplemental Fig. 9). As Ave1 is a strong contributor to virulence on plants lacking *Ve1* (de Jonge et al. 2012), selection pressure may drive the high level of conservation and lack of SNPs (Fig. 3B) in this region in race 1 strains.

Phylogenetic analyses revealed that *Ave1* was likely acquired by *V. dahliae* through horizontal gene transfer from plants (de Jonge et al. 2012). Intriguingly, however, *V. dahliae* race 1 strains do not occur as a monophyletic clade in the *V. dahliae* population (Supplemental Fig. 3A) (de Jonge et al. 2013), and thus it remains unclear how the *Ave1* locus distributed throughout the *V. dahliae* population. To assess if *Ave1* was gained or lost multiple times, we studied the genomic region surrounding the *Ave1* locus in the LS region. As expected, alignments of genome assemblies of multiple *V. dahliae* strains revealed the absence of *Ave1* in all race 2 strains (Supplemental Fig 10A). By mapping paired-end reads derived from genomic sequencing of various *V. dahliae* strains onto the genome assembly of *V. dahliae* strain JR2 we observed clear differences in coverage levels between *V. dahliae* race 1 and race 2 strains that carry or lack the *Ave1* gene, respectively (Fig. 3C; Supplemental Fig. 10B). While *V. dahliae* race 1 strains, including JR2, displayed an even level of read coverage over the *Ave1* locus, indicating that this region is similar in all race 1 stains, no read coverage of the *Ave1* gene was found for race 2 strains (Fig. 3C-D). Intriguingly, read coverage surrounding the *Ave1* gene revealed that race 2 strains can be divided into three groups depending on the exact location of the read coverage drop (Fig. 3C). Whereas one group of isolates does not display any read coverage up to 720 kb, two groups display distinct regions in which the read coverage around the *Ave1* locus drops (Fig. 3C-E: red lines around 668 kb and green lines around 672 kb). Our finding strongly suggests that the *Ave1* locus was lost multiple times from the *V. dahliae* population, pointing to strong selection pressure posed by the Ve1 immune receptor on *V. dahliae* to lose *Ave1.* In conclusion, the *Ave1* locus is situated in a highly dynamic and repeat-rich region (Fig. 3C). Therefore, the most parsimonious evolutionary scenario is that the *Ave1* locus was horizontally acquired either directly or indirectly from plants once, followed by multiple losses in independent lineages that encountered host plants that carried Ve1 or functional homologs of this immune receptor (Thomma et al. 2011; de Jonge et al. 2012; Zhang et al. 2014). It is tempting to speculate that the transposable elements that flank the *Ave1* locus (Fig. 3C) contributed to the evolution of the *Ave1* locus by facilitating swift loss of the effector gene upon recognition by the Ve1 immune receptor.

### *Lineage-specific genomic regions in* Verticillium dahliae *contain active transposable elements*

While the activity of TEs is not associated with the formation of extensive genome rearrangements, LS regions in *V. dahliae* are highly enriched for TEs, and their presence and potential activity may contribute to accelerated evolution of these genomic regions. In *V. dahliae,* the most abundant class of TEs are retrotransposons that transpose within the genome using an RNA intermediate (Wicker et al. 2007; Faino et al. 2015) (Supplemental Table 1). We assessed TE dynamics by querying the transcriptional activity of TEs using *in vitro* RNA-Seq data derived from *V. dahliae* strain JR2(de Jonge et al. 2012). Notably, the majority of TEs in *V. dahliae* are not transcribed and thus likely not active (Supplemental Table 1), while transcribed and therefore likely active TEs are found in LS regions (Supplemental Fig. 11).

To further assess if and how TEs influence the evolution of LS regions, we explored TE dynamics in the genome of *V. dahliae* strain JR2. Each copy of a TE in the genome is derived from an active ancestor that, once transposed and integrated into the genome, accumulates mutations that, over evolutionary time, will render the TE inactive. The relative age of individual TEs can thus be estimated based on sequence divergence from a consensus sequence that can be derived from present-day copies of any given TE. Using the Jukes-Cantor distance (Jukes and Cantor 1969), which corrects the divergence between TEs and their consensus sequence for multiple substitutions, we estimated the divergence times for TEs in the *V. dahliae* strain JR2 genome (Fig. 4). This analysis showed that TEs mainly transposed and expanded in two distinct periods (Fig. 4A). Notably, a considerable amount of ‘younger’ TEs, i.e. with small Jukes-Cantor distance to their consensus sequence, localizes in LS regions while the majority of ‘older’ TEs reside in the core genome (Fig. 4A). Next, we attempted to determine the relative period in which the majority of TEs in the genome of *V. dahliae* strain JR2 transposed. To this end, we derived Jukes-Cantor distributions for orthologous genes of *V. dahliae* strain JR2 and individual closely related fungi as a proxy of divergence between these species (Fig. 4C). These distributions display the same pattern of species divergence when derived from phylogenetic analyses (Supplemental Fig. 12) as well as from the Ks distributions between orthologous gene pairs (Fig. 2D). By comparing the Jukes-Cantor distributions derived from TEs and from orthologous genes, we revealed that ‘older’ TEs transposed around the separation of *V. dahliae* and *V. alfalfa,* while the ‘younger’ TEs transposed after *V. dahliae* speciation (Fig. 4C). Notably, ‘younger’ TEs tend to be transcriptionally active, while ‘older’, more diverged TEs tend to be transcriptionally silent (Fig. 4B). Thus, the expansion of younger TEs is recent and mainly concerns the active TEs localized at LS regions (Fig. 4), strongly suggesting that TE-transpositions contribute to the genetic plasticity of LS regions.

## Discussion

Many plant pathogens contain a so-called ‘two-speed’ genome where effector genes reside in genomic compartments that are considerably more plastic than the core of the genome, facilitating the swift evolution of effector catalogs that are required to be competitive in the host-pathogen arms race (Raffaele and Kamoun 2012; Dong et al. 2015). Generally, effector compartments are enriched in transposable elements (TEs), and it has been speculated that they promote genomic flexibility and drive accelerated evolution of these genomic compartments (Gijzen 2009; Haas et al. 2009; Ma et al. 2010; Raffaele et al. 2010; Rouxel et al. 2011; Raffaele and Kamoun 2012; de Jonge et al. 2013; Wicker et al. 2013; Grandaubert et al. 2014; Seidl and Thomma 2014; van Hooff et al. 2014; Dong et al. 2015; Faino et al. 2015; Seidl et al. 2015). Nevertheless, the exact role of TEs in the evolution of effector genes, as well as the molecular mechanisms and the evolutionary trajectories that govern the formation of the ‘two-speed’ genome, and the corresponding local genome plasticity, remained unknown (Raffaele and Kamoun 2012; Dong et al. 2015). Here, we studied the evolution of the ‘two-speed’ genome of the vascular wilt pathogen *V. dahliae,* which was significantly facilitated by the recent establishment of gapless whole-genome assemblies of two highly similar *V. dahliae* strains that, despite their high (>99%) degree of sequence identity, display severe genomic rearrangements (Faino et al. 2015; Thomma et al. 2015). We identified ∽2 Mb repeat-rich, lineage-specific (LS) regions between the two *V. dahliae* strains (Fig. 1; Supplemental Fig. 6-7) that are significantly enriched in TEs and contain all thus far functionally analyzed effector genes, including *Ave1* (Fig. 3C) (de Jonge et al. 2012; de Jonge et al. 2013). Moreover, we also determined a significant number of synteny breakpoints that are associated with genomic rearrangements to high resolution, and we were able to reconstitute most of the rearrangements in detail (Fig. 1; Table 1; Supplemental Fig. 5). Previously, we showed a strong association between LS compartments and the occurrence of genomic rearrangements (de Jonge et al. 2013). Even though this correlation was overestimated due to significant errors in the publically available genome assembly of *V. dahliae* strain VdLs17 (Klosterman et al. 2011) for which we later revealed a considerable number of erroneous chromosomal inversions (Faino et al. 2015) (Table 1), we nevertheless observed that three out of four large-scale LS regions are associated with chromosomal rearrangements (Fig. 1), corroborating that chromosomal rearrangements significantly contribute to the evolution of LS regions in *V. dahliae.* We were not able to associate every LS region with a genomic rearrangement (Fig. 1), like we were also not able to exactly reconstitute each genomic rearrangement (Fig. 1; Supplemental Fig. 5). However, it can easily be anticipated that complex rearrangements occurred in which genetic material in close proximity was lost. Furthermore, likely these rearrangements (continuously) occurred over longer evolutionary timescales (Supplemental Fig. 3; Supplemental Fig. 4), and subsequent rearrangement events may have erased ‘scars’ of previous rearrangements. Thus, even though not every genomic rearrangement leads to a new LS region, large-scale genomic alterations are the driving force for their formation in *V. dahliae.*

Genomic rearrangements can lead to a multitude of structural variations, including translocations, duplications and gene loss (Seidl and Thomma 2014). Here, we show that the LS compartments of *V. dahliae* strain JR2 evolved by frequent segmental duplications that yielded genetic material that subsequently obtained the freedom to diverge (Fig. 2B). Intriguingly, these duplications occurred recently, namely after divergence of *V. dahliae* and *V. alfalfae* (Fig. 2). However, this is not the only wave of duplications that shaped the *V. dahliae* genome, as an earlier duplication period was found that concerns duplications found in the core genome (Fig. 2). Notably, this duplication event was not only observed in *V. dahliae,* but also in related species including *V. alfalfae, V. albo-atrum, A. alcalophilium* and *C. higginsianum,* yet not in *F. oxysporum,* suggesting that this duplication occurred after the speciation of *F. oxysporum* from the last common ancestor of the more recently evolved fungi (Fig. 2). Given the high abundance of duplicated genes at a confined point in evolution, it is tempting to speculate that a single genome-wide event, for example a whole-genome duplication or hybridization, gave rise to these duplicates. Potentially, this particular duplication event determined that, despite their similar life style and physiology, *V. dahliae* evolved LS regions that are embedded within its core genome, while *F. oxysporum* evolved LS genetic material as conditionally dispensable chromosomes that can be horizontally transferred (Ma et al. 2010).

Based on our data, which indicate that TEs can be implicated in many yet not all genomic rearrangements, it is conceivable that homologous recombination using TE sequences as a substrate, rather than TE activity, is responsible for the establishment of gross rearrangements. In this manner, TEs have passively contributed to the genome evolution of *V. dahliae,* and to the extensive genomic plasticity that can be observed between *V. dahliae* strains (Fig. 1). In addition to the passive roles of TE in genome evolution, we also obtained evidence for the involvement of TE activity in the evolution of *V. dahliae.* Intriguingly, while the majority of TEs in *V. dahliae* is transcriptionally silent and thus inactive, transcriptionally active and thus actively transposing TEs were observed in LS regions (Fig. 4, Supplemental Fig. 9). Similarly, transcription and specific induction of TE activity has been observed previously in *V. dahliae* strain VdLs17 (Amyotte et al. 2012). These independent observations are further corroborated by our dating analyses suggesting that TEs that localize in LS regions are significantly ‘younger’ when compared with those residing in the core genome (Fig. 4). Active transposition can contribute to genome plasticity by causing gene deletions at their target sites. Moreover, TEs in proximity of genes can also profoundly influence their expression, as shown for the barley smut fungus *Ustilago hordei* where a TE insertion in the promoter of the *UhAvr1* effector changed its expression, leading to loss of host recognition (Ali et al. 2014). Interestingly, frequent gene loss is observed within the segmental duplications that constitute the LS regions (Fig. 2; Supplemental Fig. 8). Similarly, the LS effector *Ave1* is located in an LS region (Fig. 3) with several transposable elements (TE) in its direct vicinity (de Jonge et al. 2012) (Fig. 3C), and clear evidence for repeated loss in the *V. dahliae* population was obtained in our study. Although based on our data these transposons cannot directly be implicated in the acquisition of *Ave1* from plants, they likely facilitated the multiple losses of *Ave1* that are observed in independent lineages incited by selection pressure posed by the *Ve1* resistance gene of tomato, or functional homologs that may occur in other plants (Fradin et al. 2009; Thomma et al. 2011; de Jonge et al. 2012; Zhang et al. 2013). However, although *Ave1* resides in a LS region, and LS regions are characterized by the presence of active TEs, it remains unclear whether the frequent loss of *Ave1* is actually mediated by TE activity. It has similarly been hypothesized that ectopic genomic rearrangements caused by homology-based DNA repair pathways, possibly passively mediated by TEs, drive the frequent loss of the *Avr-Pita* effector gene in isolates of the rice blast fungus *Magnaporthe oryzae* that encountered the *Pita* resistance gene of rice (Orbach et al. 2000; Chuma et al. 2011). Similar processes likely contribute to the frequent recovery of *Avr-Pita* in *M. oryzae* strains relieved from selection pressure of Pita, leading to translocation and duplications of this effector gene in dynamic genomic regions (Chuma et al. 2011).

**Figure 2.**
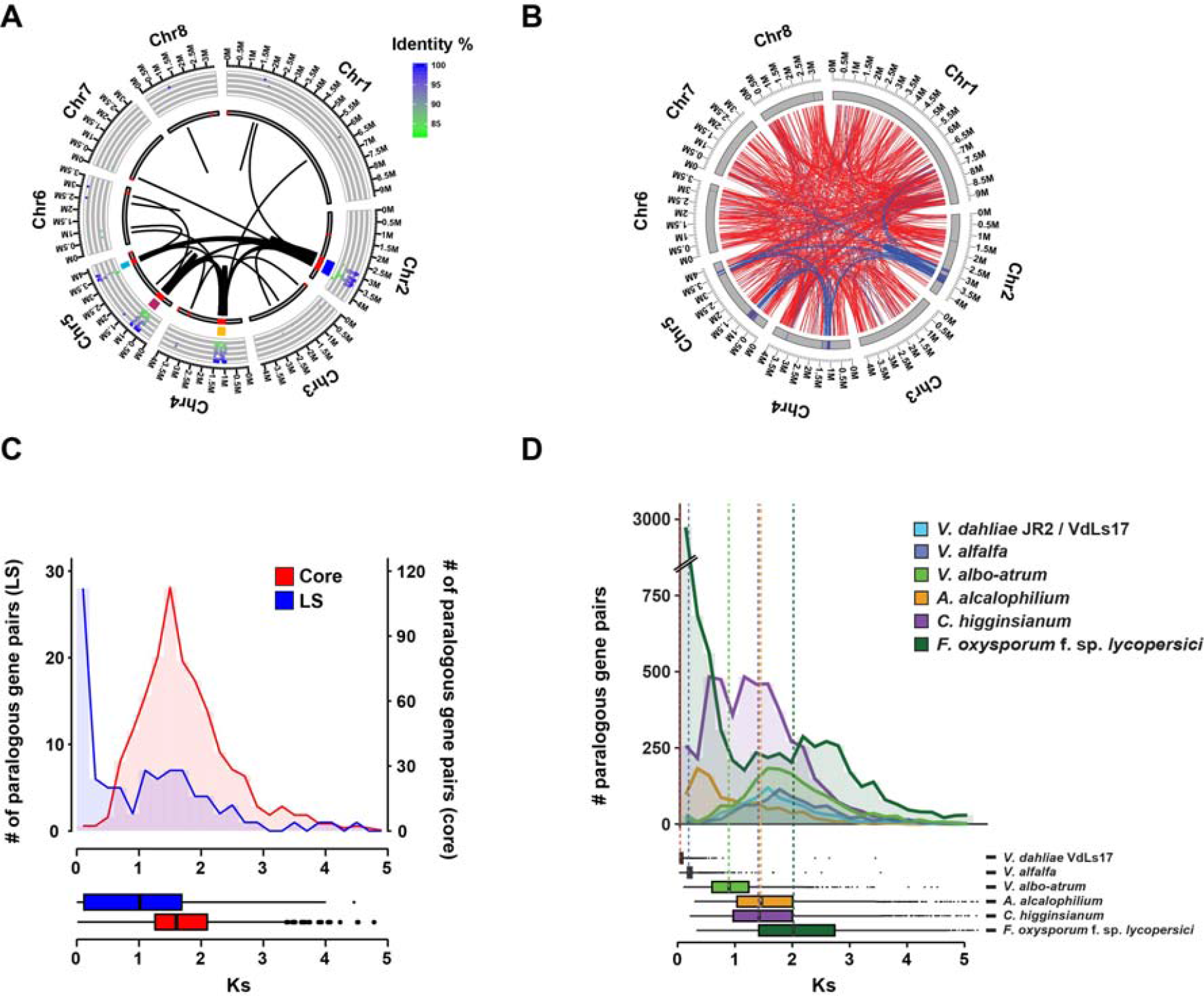
Whole-genome alignments of *Verticillium dahliae* strain JR2 reveals two duplication events. (A) Circos diagram illustrating sequence alignments within *V. dahliae* strain JR2. Black lines indicate genomic regions with sequence similarity. The inner circle shows LS regions (red lines), the middle circle indicates clusters of LS regions and the outer circle shows the identity between pairs of secondary alignments. Each cluster of LS region is colour coded: LS1 in blue, LS2 in yellow, LS3 in brown and LS4 in light blue (see Table S2). (B) Circos diagram illustrating paralogous gene pairs in *V. dahliae* strain JR2. Paralogous gene pairs of which at least one member is located at the LS regions are connected with blue lines, while paralogous gene pairs located in the core genome are connected with red lines. (C) Ks distribution of paralogs of which both genes are located in the core genome (red) or at least one paralogs is located in an LS region (blue). (D) Duplication events are estimated by calculating the Ks value for paralogous gene pairs and displayed in the line graph. Speciation events are estimated by calculating the Ks value for orthologous gene pairs based on genes from *V. dahliae* strains JR2 and their respective orthologs in the other genomes and displayed in the box plot. Distributions and median divergence times between orthologous pairs, displayed by box plots, were used to estimate relative speciation events.

Genome-wide studies in several fungi aiming to study chromatin - the complex of DNA and proteins - revealed that TE-rich regions are generally associated to highly condensed chromatin, the so-called heterochromatin, that restricts transcription and TE activity (Lewis et al. 2009; Connolly et al. 2013; Galazka and Freitag 2014). Therefore, TEs can influence the expression of neighboring genes such as effectors, as they can direct the formation of heterochromatic regions (Lewis et al. 2009). In the saprophytic fungus *Neurospora crassa,* heterochromatin formation at TEs is directed by remnants of Repeat Induced Point mutations (RIP), a pre-meiotic process that actively induces point mutations in TEs (Selker 1990; Lewis et al. 2009; Lewis et al. 2010). In the pathogenic fungus *Leptosphaeria maculans,* effectors are located in TE-rich regions that were subjected to extensive RIP mutations (Rouxel et al. 2011). Notably, effectors within these regions display signatures of nucleotide mutations caused by the RIP process, fostering rapid effector diversification (Fudal et al. 2009; Daverdin et al. 2012). LS regions in *V. dahliae* are TE-rich and effector genes are directly flanked by TEs (de Jonge et al. 2012; de Jonge et al. 2013; Seidl et al. 2015)(Fig. 3). Only few transposable elements in *V. dahliae* display typical RIP mutations, which might indicate that *V. dahliae* possessed an active RIP mechanism (Klosterman et al. 2011; Amyotte et al. 2012), likely connected to sexual cycle, in its evolutionary past. However, all currently known *V. dahliae* strains, but also other *Verticillium* species have been described as asexual and therefore RIP is no longer likely to occur. *V. dahliae* effectors in LS regions do not display evidence for RIP mutations that could contribute to effector diversification or to formation of heterochromatic regions. Despite the fact that LS regions do not show overrepresentation of secreted genes, we previously observed that effector genes residing in the LS regions are considerably overrepresented in the *V. dahliae* transcriptome upon plant infection (de Jonge et al. 2013). In the present study, we highlight that TEs that reside in LS regions are relatively young and several are transcriptionally active. Therefore, we hypothesize that these regions, in contrast to the core genomic compartments, are either not yet targeted by heterochromatin formation or carry different chromatin marks, or that LS regions represent facultative heterochromatin that can dynamically change its conformation, thereby influencing DNA accessibility and thus effector gene expression. For example, in the soybean pathogen *Phytophthora sojae* expression of the effector *Avr3a* is repressed by chromatin-based mechanism (Qutob et al. 2013; Gijzen et al. 2014). Suppression of effector expression leads to avoidance of recognition by the host surveillance system, yet allows for swift effector recovery and activation on susceptible hosts (Gijzen 2009; Qutob et al. 2013). Chromatin-based regulation of effector genes and genes encoding other virulence factors has been observed in several pathogenic fungi (Connolly et al. 2013; Qutob et al. 2013; Chujo and Scott 2014; Soyer et al. 2014; Soyer et al. 2015). This suggests that facultative heterochromatin located in dynamic compartments of the two-speed genome, e.g. LS regions in *V. dahliae,* is pivotal to regulate effector and virulence gene expression.

**Figure 3.**
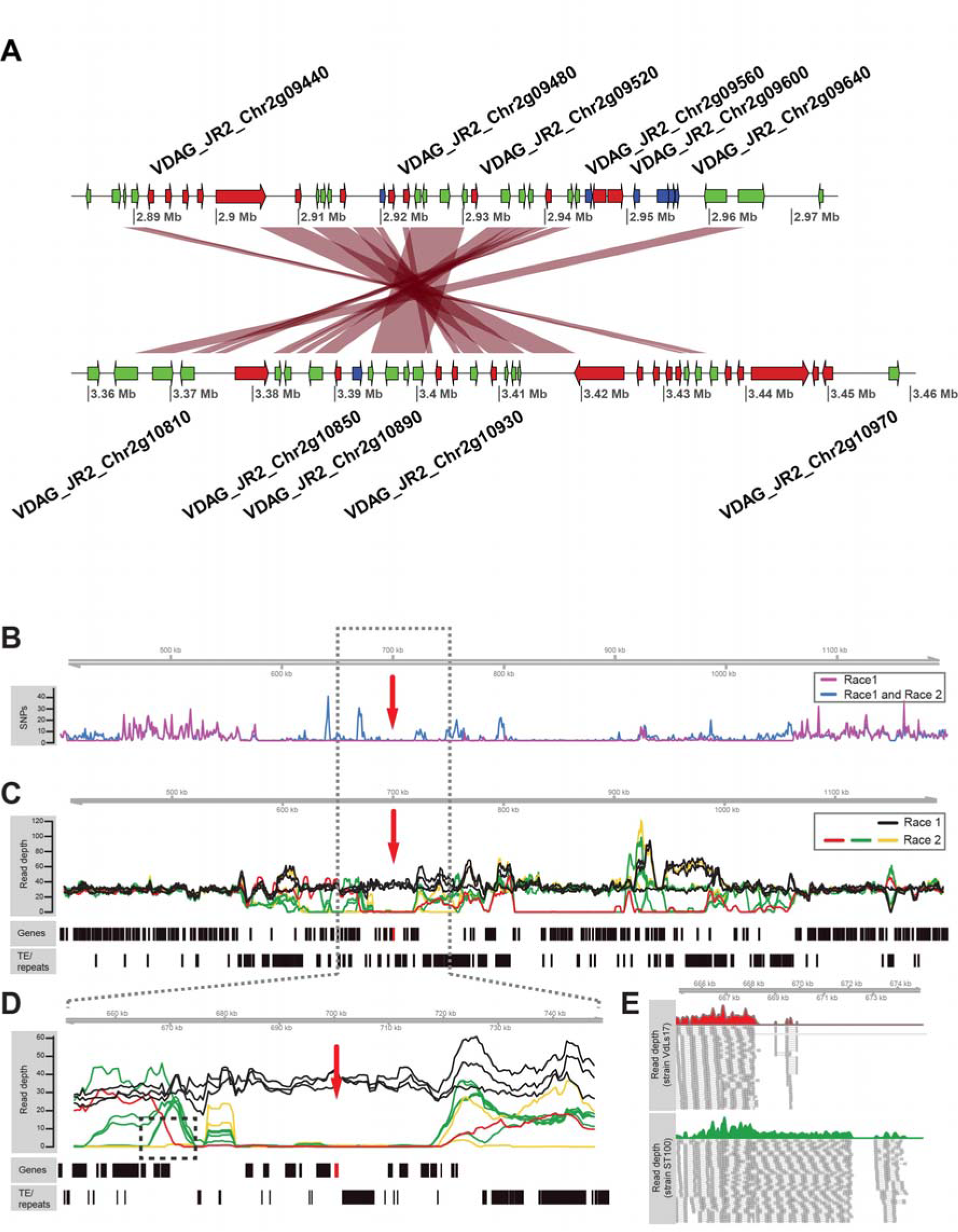
Gene loss after segmental duplications within the *V. dahliae* strain JR2 genome. (A) Example of a segmental duplication between LS regions located on chromosome 2. Red ribbons indicate regions of homology between the two loci. Blue arrows indicate gene models present only at one of the two loci while green and red arrows indicate common genes and transposable elements, respectively. (B) Single nucleotide polymorphism (SNP) density (mean number of SNPs per 1 kb) over the *Ave1* locus indicates depletion of SNPs in the *Ave1* region when compared with neighboring regions. (C) A large genomic region on chromosome 5 of *V. dahliae* strain JR2 containing the *Ave1* gene (red box) is characterized by presence/absence polymorphisms between strains. Lines indicate the corrected average read depth (per 5 kb window, 500 bp slide) of paired-end reads derived from genomic sequencing of eleven *V. dahliae* strains. Different colors indicate distinct patterns of coverage across the *Ave1* locus. Genes and transposable elements/repeats (excluding simple repeats) are indicated. (D) Magnification in of the *Ave1* locus, highlighting the four dominating coverage patterns. (E) Detailed view of paired-end reads derived from *V. dahliae* strains VdLs17 and St100, revealing distinct regions where the sequence coverage drops.

**Figure 4.**
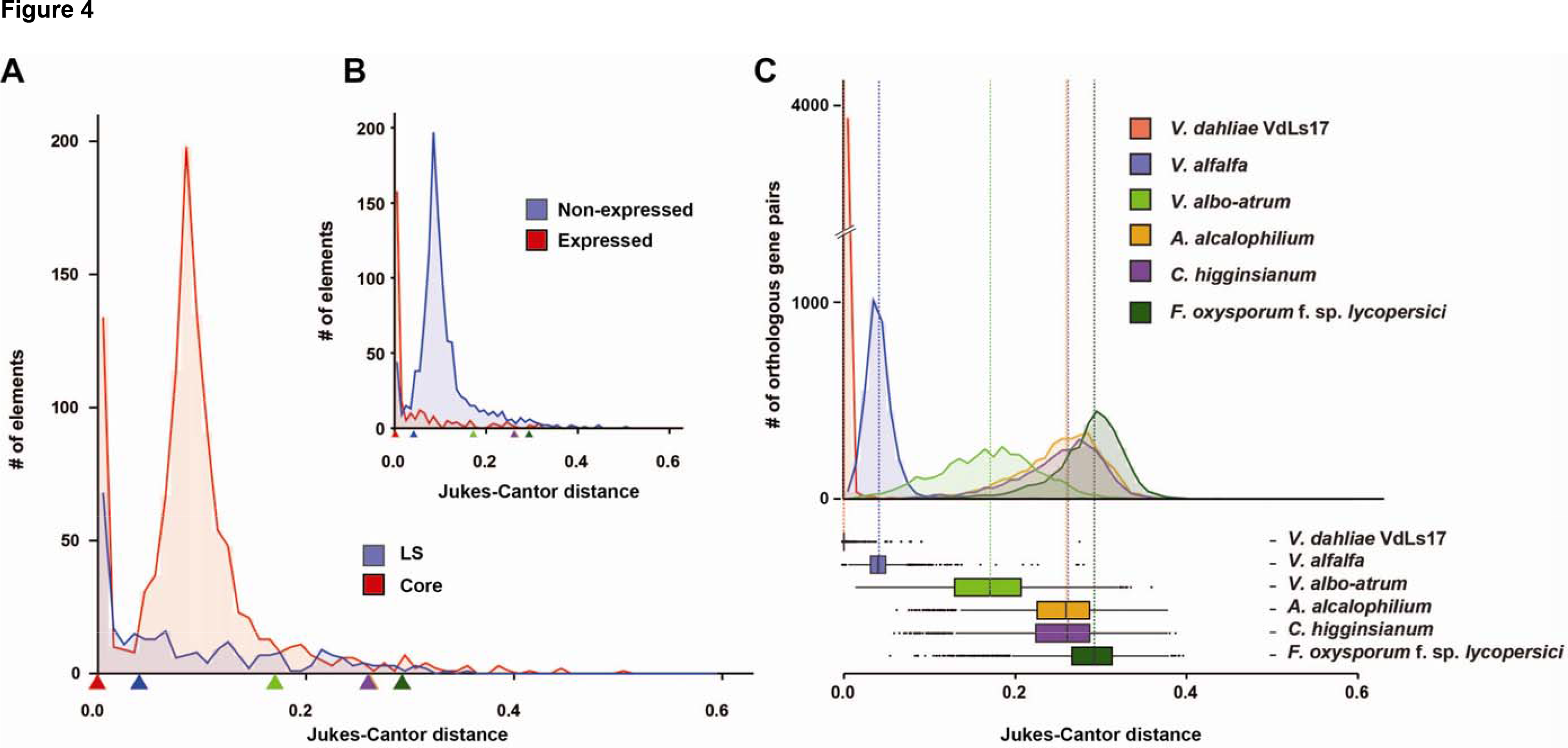
Dynamics of transposable elements in the genome of *Verticillium dahliae* strain JR2. (A) The divergence time of transposable elements identified in the genome of *V. dahliae* strain JR2 (Faino et al. 2015) was estimated using the Jukes-Cantor distance calculated between repeat copies and their consensus sequence. The distributions of divergence times between transposable elements located in the core genome (red) and in the LS regions (blue) differ. Estimations of speciation events in the evolutionary history of *V. dahliae* are indicated by triangles based on analyses in (C). (B) The distributions of divergence times between expressed/active (log_10_ (RPKM+1) > 1) transposable elements (red) and non-expressed (blue) transposable elements differ. Estimations of speciation events are indicated by triangles. (C) Speciation evens are estimated by calculating the Jukes-Cantor distance for orthologous gene pairs based on genes from *V. dahliae* strains JR2 and their respective orthologs in the other genomes. Distributions and median divergence times between orthologous pairs, displayed by box plots, were used to estimate relative speciation events.

Chromosome condensation of heterochromatic regions supposedly restricts genomic rearrangements (Galazka and Freitag 2014), yet these regions in the plant pathogens *Fusarium graminearum* are enriched for genomic rearrangements (Connolly et al. 2013). In *V. dahliae,* LS regions are formed by recent duplications and harbor active TEs, suggesting that these regions are susceptible to structural variation such as genomic rearrangements. A recent study in yeasts and mammals suggests that genomic rearrangements occur between open chromatin in regions that are in close proximity within the nucleus (Berthelot et al. 2015). Moreover, this study concludes that genomic rearrangements between TEs alone are not sufficient to fully explain the rearrangements that were observed (Berthelot et al. 2015). Expression of clustered virulence genes in the human malaria pathogen *Plasmodium falciparum* is tightly regulated by chromatin and, even though these virulence clusters are located on different chromosomes, they co-localize in close proximity within the nucleus (Ay et al. 2014). Even though these co-localized clusters have been associated with condensed, and thus repressive chromatin, extensive ectopic chromosomal rearrangements between clusters have been observed (Freitas-Junior et al. 2000; Lopez-Rubio et al. 2009; Jiang et al. 2013; Ay et al. 2014), suggesting that the chromatin structure plays roles in their evolution. In fungi, however, co-localization of effector genes in the nucleus has not yet been reported. Based on initial work in other eukaryotes, we conceive that complex chromatin structures in pathogenic fungi will not only influence coordinated effector expression in LS regions, but also genomic rearrangements, thereby further linking genome and chromatin structure to genome evolution. Thus, further insight into chromatin biology of plant pathogens will be instrumental to significantly accelerate our knowledge on the evolution of the ‘two-speed’ genome, and on fungal aggressiveness.

## Methods

### Genome annotation and dynamics

Gene predictions for the whole-genome assembly of *V. dahliae* strain JR2(Faino et al. 2015) was performed using the Maker2 software (Holt and Yandell 2011). To this end, RNA-seq reads derived from different *in vitro* and *in planta* conditions(de Jonge et al. 2012) were mapped to the genome using Tophat2 (default settings) (Trapnell et al. 2009). Additionally, gene sequences derived from previous genome annotations as well as protein sequences from 35 fungal proteomes were used as additional evidence (Klosterman et al. 2011; de Jonge et al. 2013; Seidl et al. 2015).

Homology between 13 fungal species was assessed using OrthoMCL (default settings)(Li et al. 2003). Sequence similarity between proteins was established by all-vs.-all analyses using BLASTp (E-value cutoff 1e-5, soft filtering)(Altschul et al. 1990). Ks values between gene pairs, as defined by OrthoMCL families, were calculated using the Nei-Gojobori algorithm included in the KaKs_Calculator 2.0 package(Wang et al. 2010). The coding sequences of gene pairs were aligned using protein alignment as a guide. The phylogenetic tree was generated by RAxML(Stamatakis 2006) using concatenated protein sequences of 3,492 single-copy orthologs that are conserved among eight fungal species, where only a single representative species was chosen for the fungal genera *Colletotrichum* and *Fusarium* (Supplemental Fig. 12).

Repetitive elements were identified as described in Faino et al. (2015). Briefly, repetitive elements were identified using RepeatScout, LTR_Finder and LTRharvest, and the repetitive elements identified by the different software were combined (non-redundant). Repetitive elements were further classified as described by Wicker et al. (2007). Open reading frames within the transposable elements were identified by BlastN and BlastX (Camacho et al. 2009) searches against NCBI NR databases as well as by InterProScan (Jones et al. 2014). Repetitive elements that could not be classified were defined as ‘unknown’. Expression of repetitive elements was assessed based on RNA sequencing data derived from *V. dahliae* strain JR2 grown in *in vitro* media (Czapek Dox) (de Jonge et al. 2012). Single reads were mapped onto the genome assembly of *V. dahliae* strain JR2 using Tophat2 (default parameters) (Trapnell et al. 2009). Mapped reads were summarized using the R package GenomicAlignments (‘summarizeOverlaps’) (Lawrence et al. 2013), and the expression per repetitive element, excluding simple repeats and repeats overlapping genes, as well as per gene was reported as Reads Per Kilobase of transcript per Million mapped reads (RPKM).

To estimate divergence time of transposable elements, each individual copy of a transposable element was aligned to the consensus of its family using needle, which is part of the EMBOSS package (Rice et al. 2000). The consensus sequence for each transposable element family was determined by performing multiple-sequence alignment of all copies belonging to the same family using mafft (Katoh and Standley 2013) (each individual copy needed to be longer than 400 bp). For the consensus sequence, only columns with > 1 aligned sequence (excluding gaps) were considered, for which the nucleotide occurring in the majority of sequences was used for the consensus sequence (ties, a nucleotide randomly chosen from the tie was picked). The sequence divergence between transposable elements and the consensus was corrected using the Jukes-Cantor distance, which corrects the divergence (p) by the formula d=-3/4log_e_(1-4/3p)(Jukes and Cantor 1969).

### Identification of genomic rearrangements

Whole-genome alignments between chromosomes of the genome assemblies of *V. dahliae* strains JR2 and VdLS17(Faino et al. 2015) were performed using nucmer (default settings), which is part of the MUMmer 3.0 package(Kurtz et al. 2004). To remove spurious hits, the alignments were subsequently filtered by length, retaining alignments > 15 kb and 99% identity. These parameters were chosen based on the average nucleotide identity between the two *V. dahliae* strains (99.98%), as well as the average length of unique sequences in the genome. These whole-genome alignments were further mined for genomic rearrangements and associated synteny breakpoints in *V. dahliae* strain JR2. The identified synteny breakpoints were further refined by mapping PacBio long-sequencing reads derived by genomic sequencing of *V. dahliae* strain VdLS17 to the genome of *V. dahliae* strain JR2 using Blasr (default settings) (Chaisson and Tesler 2012), followed by manual refinement. GEvo (Lyons et al. 2008) was used to identify syntenic regions between *V. dahliae* strains JR2 and VdLs17, where only gene-coding regions were used as anchors between the syntenic chromosomal regions.

To assess the presence or absence of genomic rearrangements in other *V. dahliae* strains, paired-end reads derived from genome sequencing (de Jonge et al. 2012) were mapped onto the genome of *V. dahliae* strain JR2 using BWA (BWA-mem algorithm) (Li and Durbin 2010). Genomic regions surrounding the identified genomic rearrangements (± 4 kb) were visually evaluated for the quantity of concordantly and discordantly mapped reads as well as orphan reads.

### Identification and analyses of highly dynamic genomic regions

Whole-genome alignments between chromosomes of the complete genome assemblies of *V. dahliae* strains JR2 and VdLS17 (Faino et al. 2015) were performed using nucmer (settings:-maxmatch), which is part of the MUMmer 3.0 package (Kurtz et al. 2004). Alignments separated by gaps <500 bp were merged in unique and contiguous alignments. LS regions were manually defined by identifying regions accumulating alignments breaks and TEs. Additionally, genomic reads derived from ten *V. dahliae* strains (de Jonge et al. 2012) were mapped on the *V. dahliae* strain JR2 and VdLs17, respectively, using Bowtie2 (default settings) (Langmead et al. 2009). The genomic coverage was determined by BEDtools (Quinlan and Hall 2010). The phylogenetic tree of eleven different *V. dahliae* strains was generated using RealPhy (Bertels et al. 2014) using either *V. dahliae* strain JR2 or VdLs17 as a reference strain.

The presence/absence analysis of the *Ave1* locus was performed by aligning paired-end reads from DNA sequencing of eleven *V. dahliae* strains (de Jonge et al. 2012) (including JR2) to the assembled genome of *V. dahliae* strain JR2 using BWA (BWA-mem algorithm) (Li and Durbin 2010). We averaged the raw read depth per genomic position in genomic windows (window-size 5 kb; slide 500 bp (Figure 3D; E) and window-size 500 bp; slide 100 bp (Supplemental Fig 10B), respectively), and subsequently performed a G+C correction similarly as previously described (Yoon et al. 2009). Briefly, we adjusted these averaged raw read depth (ARD) based on the observed deviation of read depth for a given G+C percentage. To this end, we first determined the average ARD for G+C percentages ranging from 0-100% (by 1%). Subsequently, we corrected the ARD using the formula ARDci = ARD_i_ * (m/m_G_+_C_), where ARDc_i_ is the corrected ARD in the ith window, ARD_i_ is the ARD in the *i*th window, m is the average ARD over all windows, and mG+C is the average ARD for all windows with the same G+C percentage as the *i*th window (Yoon et al. 2009). Additionally, the genomic reads of each individual additional *V. dahliae* strain were assembled using A5 pipeline (v.20140113) (Tritt et al. 2012). The assembled genomes were aligned to the genome assembly of *V. dahliae* strain JR2 genome using nucmer (settings:-maxmatch), which is part of the MUMmer 3.0 package (Kurtz et al. 2004). Overlaps between genomic coordinates of different genome features, e.g. lineage-specific regions, genes or transposable elements, were assessed by BEDtools (v2.24.0) (Quinlan and Hall 2010) or by the R package GenomicRanges (Lawrence et al. 2013).

Single nucleotide polymorphisms (SNPs) were identified using GATK v2.8.1 (DePristo et al. 2011). Briefly, paired-end reads derived from ten *V. dahliae* strains(de Jonge et al. 2012) were mapped onto the complete genome assembly of *V. dahliae* strain JR2 (Faino et al. 2015) using BWA (BWA-mem algorithm) (Li and Durbin 2010). Using GATK v2.8.1 (DePristo et al. 2011), mapped reads were locally realigned to minimize the number of mismatches over all reads, and subsequently genomic variants (SNPs) were called using GATK’s UnifiedGenotyper (default settings; emitting threshold 20, haploid organism) and resulting variants were quality filtered (quality > 50; phred-scaled quality score for the assertion), depth > 10 and allelic frequency > 0.9). SNPs derived from different strains were summarized in non-overlapping windows of 1 kb, and the number of SNPs derived from the individual strains were averaged per window. Absence of a SNP in a particular strain was only considered if the corresponding position displayed read coverage.

## Data access

The genome assemblies of *V. dahliae* strain JR2 and VdLS17 are available from NCBI under the assembly number GCA_000400815.2 and GCA_000952015.1, respectively (Faino et al. 2015). Paired-end reads derived from genomic sequencing of *V. dahliae* strains as well as RNA sequencing reads were generated by de Jonge et al. (2012) (PRJNA169154). Genome sequences and annotation for *Fusarium oxysporum* f. sp. *lycopersici, Colletotrichum higginsianum* were retrieved from the Ensembl fungi database (http://fungi.ensembl.org/). The data for *Acremonium alcalophilum* were downloaded from JGI database, and the data for *V. alfalfa* MS102 (Klosterman et al. 2011) were obtained from the *Verticillium* database at the Broad Institute.

## Acknowledgments

Work in the laboratory of BPHJT is support by the Research Council Earth and Life Sciences (ALW) of the Netherlands Organization of Scientific Research (NWO). MFS acknowledges the receipt of a VENI grant of ALW-NWO, project number 863.15.005.

## Author Contributions

LF conceived the study, participated in its design and coordination, performed analyses, and contributed to writing of the manuscript; MFS conceived the study, participated in its design and coordination, performed analyses, and wrote the manuscript; GCMB produced biological materials used in the study; XSK and MP performed analyses; AHJW participated in the establishment of high-quality genome assemblies; BPHJT conceived the study, participated in its design and coordination, and wrote the manuscript. All authors read and approved the final manuscript.

## References

Ali S, Laurie JD, Linning R, Cervantes-Chavez JA, Gaudet D, Bakkeren G. 2014. An immunity-tiggering effector from the barley smut fungus *Ustilago hordei* resides in an Ustilaginaceae-specific cluster bearing signs of transposable element-assisted evolution. PLoS Path 10: e1004223.

Altschul SF, Gish W, Miller W, Myers EW, Lipman DJ. 1990. Basic local alignment search tool. J Mol Biol 215: 403–410.

Amyotte SG, Tan XP, Pennerman K, Jimenez-Gasco MD, Klosterman SJ, Ma LJ, Dobinson KF, Veronese P. 2012. Transposable elements in phytopathogenic *Verticillium* spp.: insights into genome evolution and inter-and intra-specific diversification. BMC Genomics 13: 314.

Ay F, Bunnik EM, Varoquaux N, Bol SM, Prudhomme J, Vert JP, Noble WS, Le Roch KG. 2014. Three-dimensional modeling of the *P. falciparum* genome during the erythrocytic cycle reveals a strong connection between genome architecture and gene expression. Genome Res 24: 974–988.

Bertels F, Silander OK, Pachkov M, Rainey PB, van Nimwegen E. 2014. Automated reconstruction of whole-genome phylogenies from short-sequence reads. Mol Biol Evol 31: 1077–1088.

Berthelot C, Muffato M, Abecassis J, Crollius HR. 2015. The 3D organization of chromatin explains evolutionary fragile genomic regions. Cell Rep 10: 1913–1924.

Camacho C, Coulouris G, Avagyan V, Ma N, Papadopoulos J, Bealer K, Madden TL. 2009. BLAST plus : architecture and applications. BMC Bioinformatics 10: 421.

Chaisson MJ, Tesler G. 2012. Mapping single molecule sequencing reads using basic local alignment with successive refinement (BLASR): application and theory. BMC Bioinformatics 13: 238.

Chujo T, Scott B. 2014. Histone H3K9 and H3K27 methylation regulates fungal alkaloid biosynthesis in a fungal endophyte-plant symbiosis. Mol Microbiol 92: 413–434.

Chuma I, Isobe C, Hotta Y, Ibaragi K, Futamata N, Kusaba M, Yoshida K, Terauchi R, Fujita Y, Nakayashiki H et al. 2011. Multiple translocation of the *AVR-Pita* effector gene among chromosomes of the rice blast fungus *Magnaporthe oryzae* and related species. PLoS Path 7: e1002147.

Connolly LR, Smith KM, Freitag M. 2013. The *Fusarium graminearum* histone H3K27 methyltransferase KMT6 regulates development and expression of secondary metabolite gene clusters. PLoS Genet 9: e1003916.

Cook DE, Mesarich CH, Thomma BPHJ. 2015. Understanding plant immunity as a surveillance system to detect invasion. Annu Rev Phytopathol 53: 541–563.

Daverdin G, Rouxel T, Gout L, Aubertot JN, Fudal I, Meyer M, Parlange F, Carpezat J, Balesdent MH. 2012. Genome structure and reproductive behaviour influence the evolutionary potential of a fungal phytopathogen. PLoS Pathog 8: e1003020.

de Jonge R, Bolton MD, Kombrink A, van den Berg GC, Yadeta KA, Thomma BPHJ. 2013. Extensive chromosomal reshuffling drives evolution of virulence in an asexual pathogen. Genome Res 23: 1271–1282.

de Jonge R, van Esse HP, Maruthachalam K, Bolton MD, Santhanam P, Saber MK, Zhang Z, Usami T, Lievens B, Subbarao KV et al. 2012. Tomato immune receptor Ve1 recognizes effector of multiple fungal pathogens uncovered by genome and RNA sequencing. Proc Natl Acad Sci USA 109: 5110–5115.

de Visser JAG, Elena SF. 2007. The evolution of sex: empirical insights into the roles of epistasis and drift. Nature Reviews Genetics 8: 139–149.

DePristo MA, Banks E, Poplin R, Garimella KV, Maguire JR, Hartl C, Philippakis AA, del Angel G, Rivas MA, Hanna M et al. 2011. A framework for variation discovery and genotyping using next-generation DNA sequencing data. Nat Genet 43: 491–498.

Dong S, Raffaele S, Kamoun S. 2015. The two-speed genomes of filamentous pathogens: waltz with plants. Curr Opin Genet Dev 35: 57–65.

Faino L, Seidl MF, Datema E, van den Berg GCM, Janssen A, Wittenberg AHJ, Thomma BPHJ. 2015. Single-Molecule Real-Time sequencing combined with optical mapping yields completely finished fungal genomes. mBIO 6: e00936–00915.

Fedorova ND, Khaldi N, Joardar VS, Maiti R, Amedeo P, Anderson MJ, Crabtree J, Silva JC, Badger JH, Albarraq A et al. 2008. Genomic islands in the pathogenic filamentous fungus *Aspergillus fumigatus*. PLoS Genet 4: e1000046.

Flot J-F, Hespeels B, Li X, Noel B, Arkhipova I, Danchin EG, Hejnol A, Henrissat B, Koszul R, Aury J-M. 2013. Genomic evidence for ameiotic evolution in the bdelloid rotifer Adineta vaga. Nature 500: 453–457.

Fradin EF, Thomma BPHJ. 2006. Physiology and molecular aspects of Verticillium wilt diseases caused by *V. dahliae* and *V. albo-atrum*. Mol Plant Pathol 7: 71–86.

Fradin EF, Zhang Z, Ayala JCJ, Castroverde CDM, Nazar RN, Robb J, Liu CM, Thomma BPHJ. 2009. Genetic dissection of Verticillium wilt resistance mediated by tomato Vel Plant Physiol 150: 320–332.

Freitas-Junior LH, Bottius E, Pirrit LA, Deitsch KW, Scheidig C, Guinet F, Nehrbass U, Wellems TE, Scherf A. 2000. Frequent ectopic recombination of virulence factor genes in telomeric chromosome clusters of *P. falciparum*. Nature 407: 1018–1022.

Fudal I, Ross S, Brun H, Besnard AL, Ermel M, Kuhn ML, Balesdent MH, Rouxel T. 2009. Repeat-induced point mutation (RIP) as an alternative mechanism of evolution toward virulence in Leptosphaeria maculans. Molecular plant-microbe interactions: MPMI 22: 932–941.

Galazka JM, Freitag M. 2014. Variability of chromosome structure in pathogenic fungi - of ‘ends and odds’. Curr Opin Microbiol 20: 19–26.

Gijzen M. 2009. Runaway repeats force expansion of the *Phytophthora infestans* genome. Genome Biology 10: 241.

Gijzen M, Ishmael C, Shrestha SD. 2014. Epigenetic control of effectors in plant pathogens. Front Plant Sci 5.

Goodwin SB, M’Barek SB, Dhillon B, Wittenberg AH, Crane CF, Hane JK, Foster AJ, Van der Lee TA, Grimwood J, Aerts A et al. 2011. Finished genome of the fungal wheat pathogen *Mycosphaerella graminicola* reveals dispensome structure, chromosome plasticity, and stealth pathogenesis. PLoS Genet 7: e1002070.

Grandaubert J, Lowe RG, Soyer JL, Schoch CL, Van de Wouw AP, Fudal I, Robbertse B, Lapalu N, Links MG, Ollivier B et al. 2014. Transposable element-assisted evolution and adaptation to host plant within the *Leptosphaeria maculans-Leptosphaeria biglobosa* species complex of fungal pathogens. BMC Genomics 15: 891.

Haas BJ, Kamoun S, Zody MC, Jiang RH, Handsaker RE, Cano LM, Grabherr M, Kodira CD, Raffaele S, Torto-Alalibo T et al. 2009. Genome sequence and analysis of the Irish potato famine pathogen *Phytophthora infestans*. Nature 461: 393–398.

Hedges DJ, Deininger PL. 2007. Inviting instability: Transposable elements, double-strand breaks, and the maintenance of genome integrity. Mutat Res 616: 46–59.

Heitman J. 2010. Evolution of eukaryotic microbial pathogens via covert sexual reproduction. Cell host & microbe 8: 86–99.

Heitman J, Kronstad JW, Taylor JW, Casselton L. 2007. Sex in Fungi: Molecular Determination and Evolutionary Implications. ASM Press.

Holt C, Yandell M. 2011. MAKER2: an annotation pipeline and genome-database management tool for second-generation genome projects. BMC Bioinformatics 12: 491.

Inderbitzin P, Subbarao KV. 2014. *Verticillium* systematics and evolution: how confusion impedes Verticillium wilt management and how to resolve it. Phytopathology 104: 564–574.

Jiang LB, Mu JB, Zhang QF, Ni T, Srinivasan P, Rayavara K, Yang WJ, Turner L, Lavstsen T, Theander TG et al. 2013. PfSETvs methylation of histone H3K36 represses virulence genes in *Plasmodium falciparum*. Nature 499: 223–227.

Jones P, Binns D, Chang HY, Fraser M, Li W, McAnulla C, McWilliam H, Maslen J, Mitchell A, Nuka G et al. 2014. InterProScan 5: genome-scale protein function classification. Bioinformatics 30: 1236–1240.

Jukes TH, Cantor CR. 1969. Evolution of protein molecules. Mammalian protein metabolism 3: 21–132.

Katoh K, Standley DM. 2013. MAFFT multiple sequence alignment software version 7: improvements in performance and usability. Mol Biol Evol 30: 772–780.

Klosterman SJ, Atallah ZK, Vallad GE, Subbarao KV. 2009. Diversity, pathogenicity and management of *Verticillium* species. Annu Rev Phytopathol 47: 39–62.

Klosterman SJ, Subbarao KV, Kang S, Veronese P, Gold SE, Thomma BPHJ, Chen Z, Henrissat B, Lee YH, Park J et al. 2011. Comparative genomics yields insights into niche adaptation of plant vascular wilt pathogens. PLoS Path 7: e1002137.

Krejci L, Altmannova V, Spirek M, Zhao X. 2012. Homologous recombination and its regulation. Nucleic Acids Res 40: 5795–5818.

Kurtz S, Phillippy A, Delcher AL, Smoot M, Shumway M, Antonescu C, Salzberg SL. 2004. Versatile and open software for comparing large genomes. Genome Biology 5: R12.

Langmead B, Trapnell C, Pop M, Salzberg SL. 2009. Ultrafast and memory-efficient alignment of short DNA sequences to the human genome. Genome Biology 10: R25.

Lawrence M, Huber W, Pages H, Aboyoun P, Carlson M, Gentleman R, Morgan MT, Carey VJ. 2013. Software for computing and annotating genomic ranges. PLoS Comp Biol 9: e1003118.

Lewis ZA, Adhvaryu KK, Honda S, Shiver AL, Knip M, Sack R, Selker EU. 2010. DNA methylation and normal chromosome behavior in Neurospora depend on five components of a histone methyltransferase complex, DCDC. PLoS Genet 6: e1001196.

Lewis ZA, Honda S, Khlafallah TK, Jeffress JK, Freitag M, Mohn F, Schubeler D, Selker EU. 2009. Relics of repeat-induced point mutation direct heterochromatin formation in *Neurospora crassa*. Genome Res 19: 427–437.

Li H, Durbin R. 2010. Fast and accurate long-read alignment with Burrows-Wheeler transform. Bioinformatics 26: 589–595.

Li L, Stoeckert CJ, Jr., Roos DS. 2003. OrthoMCL: identification of ortholog groups for eukaryotic genomes. Genome Res 13: 2178–2189.

Lopez-Rubio J-J, Mancio-Silva L, Scherf A. 2009. Genome-wide analysis of heterochromatin associates clonally variant gene regulation with perinuclear repressive centers in malaria parasites. Cell host & microbe 5: 179–190.

Lyons E, Pedersen B, Kane J, Alam M, Ming R, Tang HB, Wang XY, Bowers J, Paterson A, Lisch D et al. 2008. Finding and Comparing Syntenic Regions among *Arabidopsis* and the Outgroups Papaya, Poplar, and Grape: CoGe with Rosids. Plant Physiol 148: 1772–1781.

Ma LJ, van der Does HC, Borkovich KA, Coleman JJ, Daboussi MJ, Di Pietro A, Dufresne M, Freitag M, Grabherr M, Henrissat B et al. 2010. Comparative genomics reveals mobile pathogenicity chromosomes in *Fusarium*. Nature 464: 367–373.

McDonald BA, Linde C. 2002. Pathogen population genetics, evolutionary potential, and durable resistance. Annu Rev Phytopathol 40: 349–379.

Orbach MJ, Farrall L, Sweigard JA, Chumley FG, Valent B. 2000. A telomeric avirulence gene determines efficacy for the rice blast resistance gene Pi-ta. Plant Cell 12: 2019–2032.

Quinlan AR, Hall IM. 2010. BEDTools: a flexible suite of utilities for comparing genomic features. Bioinformatics 26: 841–842.

Qutob D, Chapman BP, Gijzen M. 2013. Transgenerational gene silencing causes gain of virulence in a plant pathogen. Nature Commun 4: 1349.

Raffaele S, Farrer RA, Cano LM, Studholme DJ, MacLean D, Thines M, Jiang RH, Zody MC, Kunjeti SG, Donofrio NM et al. 2010. Genome evolution following host jumps in the Irish potato famine pathogen lineage. Science 330: 1540–1543.

Raffaele S, Kamoun S. 2012. Genome evolution in filamentous plant pathogens: why bigger can be better. Nature reviews Microbiology 10: 417–430.

Rice P, Longden I, Bleasby A. 2000. EMBOSS: The European molecular biology open software suite. Trends Genet 16: 276–277.

Rouxel T, Grandaubert J, Hane JK, Hoede C, van de Wouw AP, Couloux A, Dominguez V, Anthouard V, Bally P, Bourras S et al. 2011. Effector diversification within compartments of the *Leptosphaeria maculans* genome affected by Repeat-Induced Point mutations. Nature Commun 2: 202.

Rovenich H, Boshoven JC, Thomma BPHJ. 2014. Filamentous pathogen effector functions: of pathogens, hosts and microbiomes. Curr Opin Plant Biol 20C: 96–103.

Seidl MF, Faino L, Shi-Kunne X, van den Berg GC, Bolton MD, Thomma BPHJ. 2015. The genome of the saprophytic fungus *Verticillium tricorpus* reveals a complex effector repertoire resembling that of its pathogenic relatives. Mol Plant-Microbe Interact 28: 362–373.

Seidl MF, Thomma BPHJ. 2014. Sex or no sex: evolutionary adaptation occurs regardless. Bioessays 36: 335–345.

Selker EU. 1990. Premeiotic instability of repeated sequences in Neurospora crassa. Annu Rev Genet 24: 579–613.

Soyer JL, El Ghalid M, Glaser N, Ollivier B, Linglin J, Grandaubert J, Balesdent MH, Connolly LR, Freitag M, Rouxel T et al. 2014. Epigenetic control of effector gene expression in the plant pathogenic fungus *Leptosphaeria maculans*. PLoS Genet 10: e1004227.

Soyer JL, Rouxel T, Fudal I. 2015. Chromatin-based control of effector gene expression in plant-associated fungi. Curr Opin Plant Biol 26: 51–56.

Speijer D, Lukes J, Elias M. 2015. Sex is a ubiquitous, ancient, and inherent attribute of eukaryotic life. Proc Natl Acad Sci USA 112: 8827–8834.

Stamatakis A. 2006. RAxML-VI-HPC: maximum likelihood-based phylogenetic analyses with thousands of taxa and mixed models. Bioinformatics 22: 2688–2690.

Thomma BPHJ, Nurnberger T, Joosten MH. 2011. Of PAMPs and effectors: the blurred PTI-ETI dichotomy. Plant Cell 23: 4–15.

Thomma BPHJ, Seidl MF, Shi-Kunne X, Cook DE, Bolton MD, van Kan JA, Faino L. 2015. Mind the gap; seven reasons to close fragmented genome assemblies. Fungal Genet Biol doi:10.1016/j.fgb.2015.08.010.

Thon MR, Pan H, Diener S, Papalas J, Taro A, Mitchell TK, Dean RA. 2006. The role of transposable element clusters in genome evolution and loss of synteny in the rice blast fungus Magnaporthe oryzae. Genome biology 7: R16.

Trapnell C, Pachter L, Salzberg SL. 2009. TopHat: discovering splice junctions with RNA-Seq. Bioinformatics 25: 1105–1111.

Tritt A, Eisen JA, Facciotti MT, Darling AE. 2012. An integrated pipeline for de novo assembly of microbial genomes. PloS one 7: e42304.

van Hooff JJ, Snel B, Seidl MF. 2014. Small homologous blocks in *Phytophthora* genomes do not point to an ancient whole-genome duplication. Genome Biol Evol 6: 1079–1085.

Wang D, Zhang Y, Zhang Z, Zhu J, Yu J. 2010. KaKs_Calculator 2.0: a toolkit incorporating gamma-series methods and sliding window strategies. Genomics Proteomics Bioinformatics 8: 77–80.

Wicker T, Oberhaensli S, Parlange F, Buchmann JP, Shatalina M, Roffler S, Ben-David R, Dolezel J, Simkova H, Schulze-Lefert P et al. 2013. The wheat powdery mildew genome shows the unique evolution of an obligate biotroph. Nat Genet 45: 1092–1096.

Wicker T, Sabot F, Hua-Van A, Bennetzen JL, Capy P, Chalhoub B, Flavell A, Leroy P, Morgante M, Panaud O et al. 2007. A unified classification system for eukaryotic transposable elements. Nature Reviews Genetics 8: 973–982.

Yoon ST, Xuan ZY, Makarov V, Ye K, Sebat J. 2009. Sensitive and accurate detection of copy number variants using read depth of coverage. Genome Res 19: 1586–1592.

Zhang Z, Esse HP, Damme M, Fradin EF, Liu CM, Thomma BPHJ. 2013. Ve1 mediated resistance against *Verticillium* does not involve a hypersensitive response in *Arabidopsis*. Mol Plant Pathol 14: 719–727.

Zhang Z, Song Y, Liu CM, Thomma BPHJ. 2014. Mutational analysis of the Ve1 immune receptor that mediates *Verticillium* resistance in tomato. PloS one 9: e99511.

